# Novel Algorithms for the Taxonomic Classification of Metagenomic Linked-Reads

**DOI:** 10.1101/549667

**Authors:** David C. Danko, Dmitry Meleshko, Daniela Bezdan, Christopher Mason, Iman Hajirasouliha

## Abstract

We present KrakenLinked, a metagenomic read classifier for Linked-Reads. We have formulated two algorithms for read classification of metagenomic samples using linked reads: tree pruning and taxa promotion. Tree pruning improves specificity while taxa promotion improves sensitivity. Used together the algorithms improve taxonomic classification of linked-reads compared to short reads, particularly reducing false identification of taxa. We have implemented these algorithms as functions in KrakenUniq which we make available as KrakenLinked.

## Introduction

The microbiome has emerged as an important source of information in biological systems and metagenomics provides a rich source of data about the microbiome. One of the most fundamental tasks in metagenomic analysis is taxonomic classification where DNA sequences are assigned to likely taxa of origin. While a number of tools exist for taxonomic classification there remains significant room for improvement (McIntyre et al., 2017). Third generation sequencing technology presents a promising platform for future discoveries.

Long-read sequencing technologies (e.g. PacBio, Oxford Nanopore) promise the ability to make more specific taxonomic classifications in metagenomics than Next Generation Seqeuncing (NGS). However these technologies are expensive and require large amounts of input DNA which limits the resolution they can achieve in practice. An alternative is Linked-Read sequencing.

Low-cost, and low-input (*∼*1ng) DNA library preparation techniques using microfluidic methods are available from Moleculo/Illumina, and 10x Genomics. With these technologies, input DNA is sheared into long fragments of *∼*10-100kbp. After shearing, a 3’ barcode is ligated to short reads from the same long fragment of DNA. Finally, the short reads and barcodes are sequenced using standard sequencing Next Generation Sequencing platforms. This process is referred to as Linked-Read sequencing. We refer the reader to Zheng et al. (2016) for a more detailed explanation of the process of Linked-Read sequencing.

Reads with matching 3’ barcodes are referred to as a read cloud. The number of 3’ barcodes is limited in existing systems (*∼*1,000,000 for the 10x Genomics system) which causes reads from different fragments to be present in the same read cloud. In our work we observed that most read-clouds contained DNA from 2-20 long fragments. However because the total number of fragments in a read cloud is limited compared to the sample as a whole, and because it is common to have multiple reads from the same fragment, read-clouds provide significantly more information than random assortments of reads.

In general taxonomic assignment with short reads alone is limited. Since they span a relatively small stretch of DNA short reads may not contain markers that can accurately be identified at a species level even if the read is from a well characterized species. This issue is compounded for long reads. Long read technologies are more likely to contain species specific markers but high base error rates make it more difficult to identify SNPs or small indels that may distinguish strains. Even with comparatively high quality Illumina reads it can be difficult to distinguish SNPs from low abundance strains from sequencing errors (Kashtan et al., 2014).

Linked-Reads provide more information than short reads which can be potentially used for better taxonomic classification. If reads from the same long fragment of DNA are identified via assembly (e.g. Athena (Bishara et al., 2018)) or by barcode deconvolution (e.g. Minerva (Danko et al., 2019)) the markers from both reads could be used together for taxonomic classification. However, read-clouds provide useful additional taxonomic information even without clearly identifying which reads originated from the same fragment. A pair of reads in the same read-cloud are much more likely to have originated from the same genome than a randomly selected pair of reads would be.

In principle this information can be used to improve microbial assembly, taxonomic classification, and identify horizontally transferred sequences. Metagenomic samples are typically very noisy and Linked-Reads may be useful to reduce this noise. In this paper, we develop two algorithms that use Linked-Reads to improve taxonomic classification. We implement these algorithms in a modified version of the KrakenUniq classifier (Breitwieser et al., 2018; Wood and Salzberg, 2014) and demonstrate that they improve overall taxonomic classification, particularly reducing the number of false positive taxonomic identifications.

### Kraken

Kraken and KrakenUniq are fast taxonomic profilers based on identifying taxa specific kmers. KrakenUniq is an extension of Kraken that counts the number of unique kmers found at each rank. Though this feature is useful it is not strictly relevant to the work presented here.

Briefly, Kraken builds a database from a taxonomic tree and a set of reference sequences and find kmers that can be unambiguously assigned to different taxonomic ranks. A map from kmers to taxonomic assignment is retained as is the taxonomic tree itself. The RAM usage of kraken is directly related to the number of kmers in its database and typically kmers are removed uniformly at random until a database is reduced to a desired size. The total number of kmers in a database is proportion to Kraken’s sensitivity while the length of *k* is related to its specificity. By default Kraken uses a *k* of 31. A study by McIntyre et al. (2017) found Kraken to have a high false positive and low false negative rate.

To classify reads Kraken breaks the reads into kmers. The kmers are matched to the provided database, which can be accomplished in constant time via hashing. If a kmer is found in the database its taxonomic rank is saved. A taxonomic tree is constructed from all of the annotations for a read and the path with the highest weight from root to leaf is chosen as a reads classification. If multiple paths have the same weight the lowest common ancestor of the paths is returned as the classification.

KrakenUniq builds off of Kraken by approximately counting the number of marker kmers identified for each taxa using a hyperloglog counter. This information allows downstream quality evaluations without significantly affecting runtime performance. Though the enhancements in KrakenUniq are not strictly relevant to KrakenLinked we elected to modify KrakenUniq to retain the desirable downstream benefits.

### Our Contribution

We present KrakenLinked, an enhancement of Kraken and KrakenUniq, designed to improve metagenomic taxonomic classification with Linked-Reads. Like Kraken it breaks reads into kmers and tries to identify reference taxa which uniquely contain those markers. Reads are classified to the the most specific rank for which they have a marker. This leads to good sensitivity but a high false positive rate of taxonomic calls.

KrakenLinked enhances Kraken by adding two algorithms specific to Linked-Reads: tree pruning and taxa promotion. Though implemented for Kraken specifically these algorithms should apply to any marker based metagenomic classifier. Tree pruning is designed to reduce Kraken’s false positive rate, possibly at the expense of specificity while read promotion is designed to increase specificity at the risk of creating false positives. The algorithms can be used together and partly mitigate the negative effects of each other.

## Methods

We present two algorithms which can be used for taxonomic classification of Linked-Reads. Tree pruning increases specificity at the expense of sensitivity. Taxa promotion increases sensitivity but can sometimes result in misclassification. We describe how both algorithms work in general and how they are implemented in KrakenLinked. The algorithms may be used in tandem.

### Tree Pruning

Tree pruning removes markers with low abundance within a read-cloud. Pruning relies on the assumption that the taxa specific markers used by a classifier are somewhat noisy. An individual read has a limited number of markers so it is difficult to distinguish true signal from noise. However, a read-cloud is likely to have multiple reads from the same fragment making detection easier (in our first dataset read clouds with at least 8 reads averaged 3.21 reads (s.d. 2.54) from each fragment within the read-cloud). Tree pruning removes markers that may be found at low levels within individual reads but have little overall support in a read-cloud. In principle this method may result in false negatives.

Within KrakenLinked tree pruning takes a read-cloud and minimum number of markers. The minimum may be expressed as an absolute value or as a fraction of the number of reads in the read-cloud. In practice removing singletons effectively reduces false positives. A taxonomic tree is constructed from all markers in any read in the read-cloud with the weight of each node equivalent to the total number of markers in the read-cloud specific to that node. The algorithm proceeds iteratively. At each iteration each leaf node with fewer markers than the minimum is removed and the leaf’s marker count is added to its parent. The algorithm proceeds until no leaf nodes are below the minimum or only the root node remains. The pruned tree is then returned. This process is shown in figure 1.

**Figure 1:**
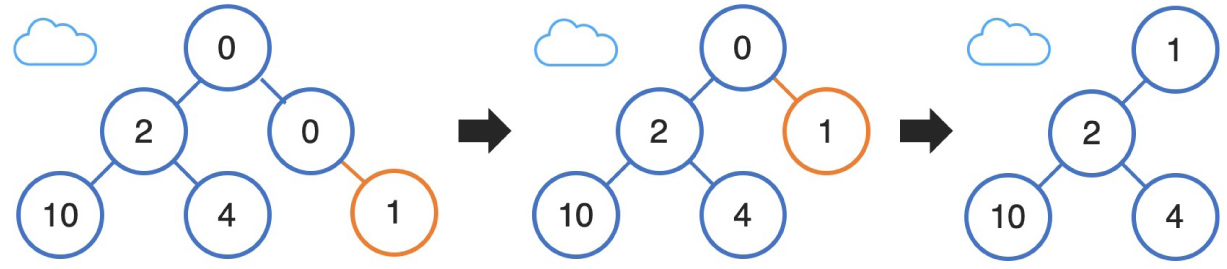
Schematic showing the tree pruning process. 1) A tree is constructed from the taxonomic assignments of all kmers in a read-cloud. 2) Low abundance leaf nodes are removed and their weight assigned to their parent. 3) This process is repeated until the tree is unchanged. The root node is never removed. Takes a single parameter: the minimum leaf-node weight.

Once a pruned tree is generated from the read-cloud each read may be individually classified to a particular taxa. A taxonomic marker tree is constructed from each read by the same method as Kraken. This tree is pruned to contain only nodes that are present in the pruned tree of the read-cloud. After the pruned read tree is constructed taxonomic assignment is found via Kraken’s typical method of finding the leaf node with the highest leaf to root weight. The highest weight path is returned as the overall taxonomic assignment of the read. Tree pruning can only make the taxonomic rank to which a read would be assigned less specific than it would be without pruning.

### Read Promotion

Read promotion increases the specificity of a read’s taxonomic assignment based on other assignments within the read-cloud. Read-clouds include DNA from a limited number of fragments, the intuition underlying read promotion is that a less specific taxonomic assignment can be assumed to be from the same organism as a more specific taxonomic assignment. Depending on the situation it may be desirable to limit the absolute number of taxonomic ranks a classification can be promoted or to set a maximum rank from which promotion may begin. Classifications are only promoted in cases where promotion is unambiguous. We originally formulated the idea of taxa promotion as a part of an earlier study (Danko et al., 2019). In that work taxa promotion was agnostic to the method used to classify reads, which potentially allowed more ambiguity than the method presented here.

In KrakenLinked read promotion works by finding the taxonomic classification of a read both with and without tree pruning. If the classifications disagree the read is considered ineligible for promotion. If the two classifications agree the classification is promoted to the most specific unambiguous rank available based on the raw (not pruned) marker tree for the entire read-cloud. A promotion is considered unambiguous if and only if there is exactly one descendant of the current rank with non-zero weight. If a rank has multiple non-zero paths it may be promoted only until the paths diverge. KrakenLinked allows the maximum number of promotions to be specified as well as the minimum starting distance from the root in order to promote a classification but these parameters made little difference in practice. This process is illustrated in figure 2.

**Figure 2:**
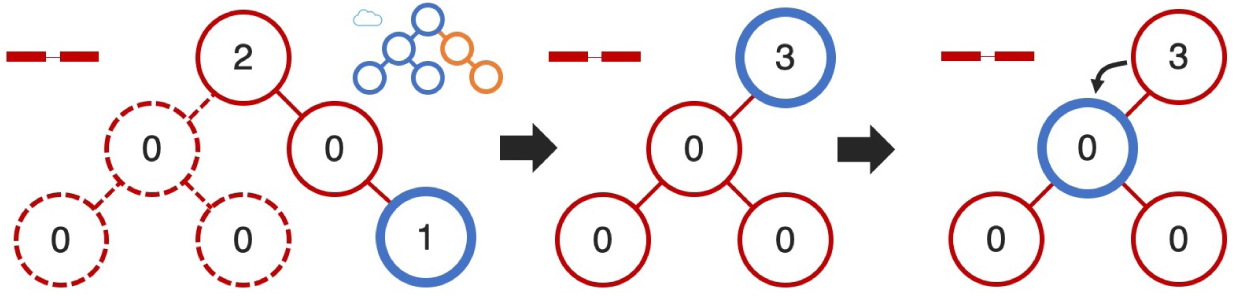
Schematic showing the process of read promotion. 1) A tree is constructed from the taxonomic assignment of all kmers in a single read. The tree is assigned the taxonomy of the root to leaf path with the highest weight (unchanged from Kraken). 2) Next the tree is pruned to contain the same nodes as the read-cloud tree. 3) The taxonomic assignment of the original tree is now promoted to the lowest non-branching child of the the original assignment or the lowest child after a defined number of hops (whichever is higher). Takes a single parameter: the maximum number of hops allowed.

## Results

### Data Sets

We tested KrakenLinked using two real data sets from two microbial mock communities originally presented in our recent study, Minerva (Danko et al. (2019)). To prepare these data sets, roughly 1ng of high molecular weight (HMW) DNA was extracted from each sample. The HMW DNA was processed using a 10x Chromium instrument and each library was sequenced on an Illumina HiSeq with 2*×*150 paired-end reads. Roughly 20M reads were generated for each sample, for testing we selected 10M reads from each while ensuring that we only selected complete barcodes. We have reproduced the description of these datasets here for convenience. The first community (Dataset 1) contained five bacterial species: *E. coli, Enterobacter cloacae, Micrococcus luteus, Pseudomonas antarctica*, and *Staph. epidermidis*. The second community (Dataset 2) contained 8 bacterial species and 2 fungi: *Bacillus subtilis, Cryptococcus neoformans, Enterococcus faecalis, E. coli, Lactobacillus fermentum, Listeria monocytogenes, Pseudomonas aeruginosa, Saccharomyces cerevisiae, Salmonella enterica*, and *Staphylococcus aureus*. The relative abundance of each species in each data set is listed in table 2.

We used to provide as realistic a dataset as possible. All species in the mock communities had well characterized genomes making taxonomic assignment easy. The mock communities chosen are standard microbial positive controls as noted by (Mason et al., 2017).

We used *Long Ranger BASIC* to attach barcodes to reads and perform error correction on barcodes (https://support.10xgenomics.com/genome-exome/software/pipelines/latest/advanced/other-pipelines). Both samples have a similar number of reads per barcode. Sample 2 had more species represented in each barcode on average, though not necessarily more fragments since fragments can originate from the same genome. Statistics about the datasets are summarized in table 1.

**Table 1:**
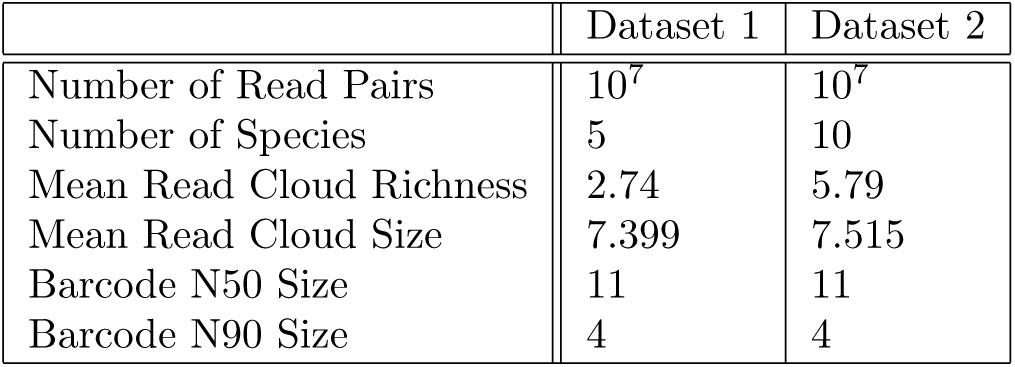
Dataset Properties

|  | Dataset 1 | Dataset 2 |
| --- | --- | --- |
| Number of Read Pairs | $10^7$ | $10^7$ |
| Number of Species | 5 | 10 |
| Mean Read Cloud Richness | 2.74 | 5.79 |
| Mean Read Cloud Size | 7.399 | 7.515 |
| Barcode N50 Size | 11 | 11 |
| Barcode N90 Size | 4 | 4 |

**Table 2:** Taxa Detail, Relative Abundance is based on read counts and is not adjusted for genome size

| Taxa | Ref. Genome Size (Mb) | Rel. Abund. Dataset 1 | Rel. Abund. Dataset 2 |
| --- | --- | --- | --- |
| *E. coli* | 5.4 | 29.39 | 31.54 |
| *Enterobacter cloacae* | 5.7 | 31.37 | n/a |
| *Micrococcus luteus* | 2.5 | 12.19 | n/a |
| *Pseudomonas antarctica* | 6.7 | 11.48 | n/a |
| *Staph. epidermidis* | 2.6 | 15.57 | n/a |
| *Bacillus subtilis* | 3.9 | n/a | 3.23 |
| *Lactobacillus fermentum* | 1.9 | n/a | 12.82 |
| *Listeria monocytogenes* | 3.0 | n/a | 3.64 |
| *Pseudomonas aeruginosa* | 6.8 | n/a | 14.70 |
| *Salmonella enterica* | 4.8 | n/a | 28.95 |
| *Staph. aureus* | 2.9 | n/a | 1.50 |
| *Enterococcus faecalis* | 3.0 | n/a | 3.50 |
| *Cryptococcus neoformans* | 18.9 | n/a | 0.05 |
| *Saccaromyces cerevisiae* | 19.1 | n/a | 0.016 |

We determined the actual genome of origin for each read by mapping reads to the source genomes using Bowtie2 (Langmead and Salzberg, 2012) (very sensitive presets). We filtered our datasets to include only read clouds with 8 reads or more (*∼*65% of all reads) and used only the first read in each pair. We used a KrakenUniq database based on all of RefSeq Microbial (ca. April 2017) including *∼*200GB of data.

### Evaluation Metrics

We use four metrics to evaluate the performance of KrakenLinked. We refer to *concordant* taxonomic assignments when the assignment is the true species (as determined by Bowtie2) an ancestor of the true species or a descendant. We refer to *exact* taxonomic assignments when the assignment is the true species or a descendant thereof.

The *Rate Correct* is the fraction of classified reads which were classified concordantly with their actual assignment. *Concordant Match* refers to the total fraction of reads (classified or not) that concordantly matched. *Richness* is the total number of species detected in the whole sample (without any secondary filtering). *Exact match* is the total fraction of reads which were classified exactly.

### Performance of Tree Pruning

In figure 3 we summarize the results of tree-pruning on Kraken’s output without taxa-promotion. Tree pruning reduces the rate at which classified reads are assigned to the wrong taxa, increases the total fraction of concordant matches, and reduces false positive species hits. However, tree-pruning slightly reduces the total fraction of reads which are correctly classified at the species rank.

**Figure 3:**
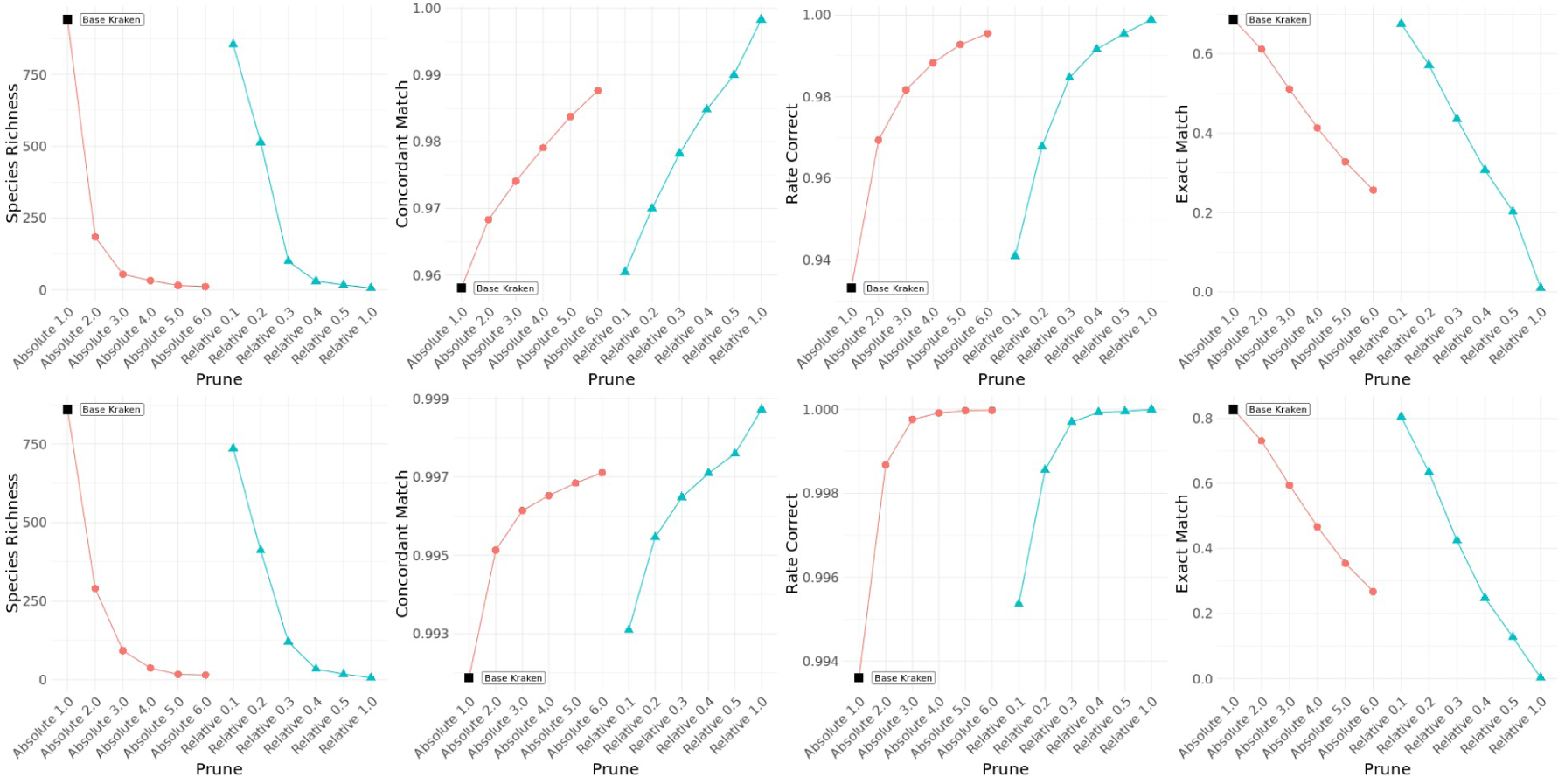
Top dataset 1, bottom dataset 2. From left to right. 1) The fraction of classified reads which were assigned to the correct taxa. 2) The fraction of reads which were classified concordantly 3) The total number of species observed in each sample (there are 5 species in the control) 4) The fraction of reads which were correctly classified at at least the species of rank

Note that tree pruning takes a single parameter, the minimum weight of a leaf-node. This parameter can be given as a constant for the whole dataset (red in figure 3) or given as a fraction of the number of reads in the read-cloud currently being processed (blue in figure 3). Increasing the minimum weight increases downstream specificity with marginally decreasing benefit but also decreases the sensitivity more or less linearly.

### Performance of Taxa Promotion

In figure 4 we summarize the results of taxa-promotion on Kraken’s output without tree-pruning. Taxa promotion increases the total fraction of reads which are correctly classified at the species rank (or more specific) but slightly increases the rate of misclassified reads.

**Figure 4:**
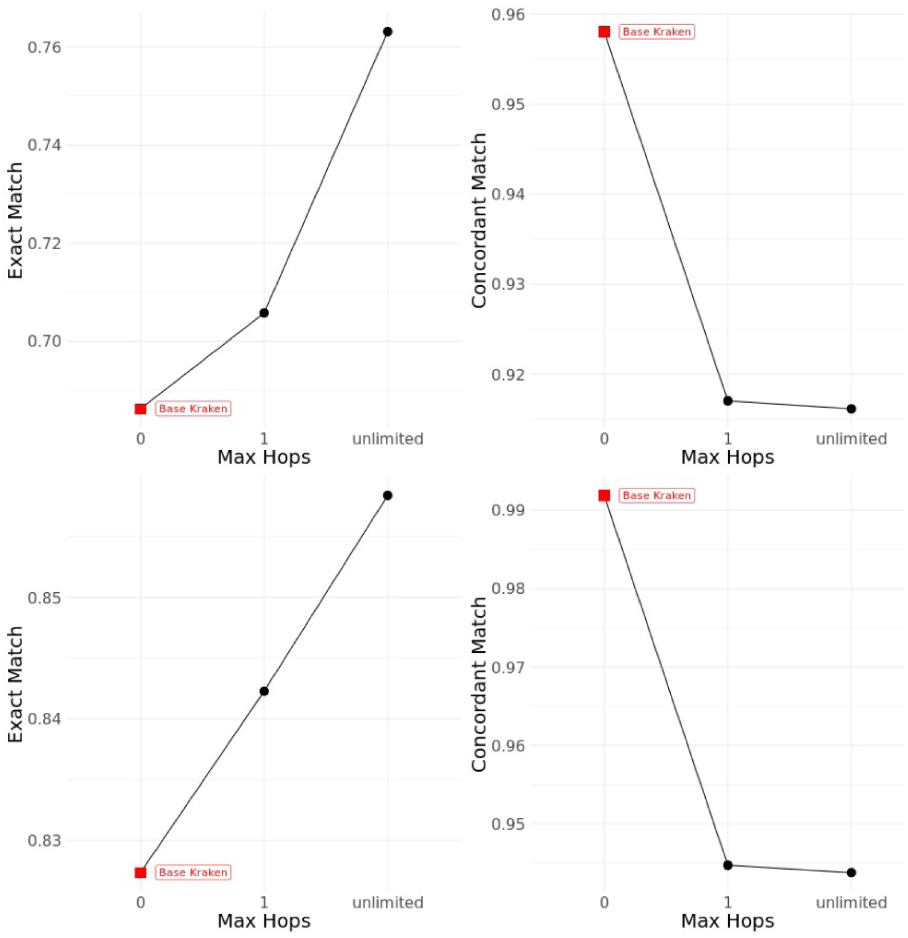
Top dataset 1, bottom dataset 2. From left to right. 1) The fraction of reads which were correctly classified at at least the species of rank 2) The fraction of reads which were classified concordantly

Taxa promotion takes one parameter, the maximum number of ranks a read may be promoted, which we term’hops’. With unlimited hops a read may be promoted from the root of the taxonomic tree to the species level (or even lower). Reads that were unclassified at any rank are never promoted. We experimented with limiting the rank from which reads could be promoted to more specific ranks (e.g. family, genus) but found this had little effect on downstream results. In general, we found that limiting the number of allowed hops was not desirable. Most of the misclassifications occurred with a single hop but failed to significantly improve the classification rank for a number of reads.

### Joint Performance of Tree Pruning and Taxa Promotion

In figure 5 we show that tree-pruning and taxonomic promotion can be used to mitigate their respective reductions to sensitivity and specificity. As shown in figure 3 aggressive tree pruning reduces the rate at which reads match at exactly the right rank. Allowing taxa promotion with unlimited hops increases the fraction of exact matches when used with tree pruning. Analogously, taxa promotion increases the misclassification rate, using tree-pruning in combination with taxa promotion mitigates this effect. Tree pruning and taxonomic promotion may be used in concert to provide a tuneable sensitivity and specificity while greatly reducing the number of false positive species calls.

**Figure 5:**
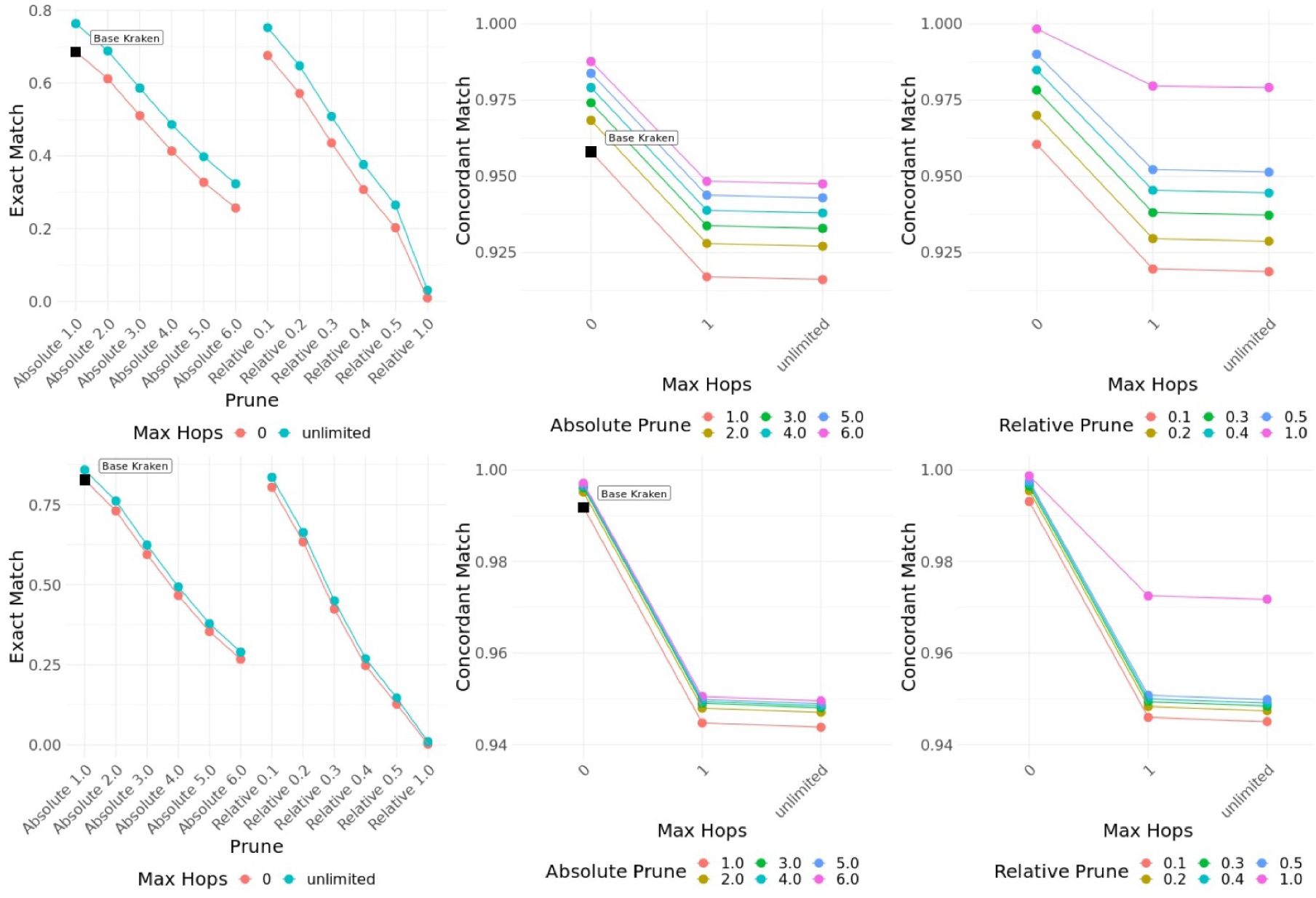
Top dataset 1, bottom dataset 2. From left to right. 1) The fraction of reads which were correctly classified at at least the species of rank 2) The fraction of reads which were classified concordantly, split for clarity.

### Runtime and Performance

RAM usage is largely unchanged from KrakenUniq, runtime is slightly slower. On dataset 1 KrakenUniq took 581 seconds, 715 seconds on dataset 2. KrakenLinked took on average of 694 seconds on dataset 1, and 963 seconds on dataset2.

## Discussion

We have introduced two algorithms to improve the taxonomic classification of marker based metagenomic classifiers using Linked-Reads and have implemented these algorithms in KrakenLinked. These algorithms, tree pruning and taxa promotion, provide adjustable sensitivity and specificity when used in concert. Compared to short reads overall results may be improved.

However, we have only evaluated KrakenLinked on two datasets containing a small number of well characterized species. This makes the classification problem tractable as essentially all of the sequence in our samples came from well known taxa but left relatively little room to improve on the results of short read classifiers. It is not immediately clear how KrakenLinked would perform on a more complex dataset.

Additionally, the algorithms presented here are based on a simple model of Linked-Reads. They do not account for the distribution of reads within a fragment nor do they take advantage of barcode deconvolution. We believe it would be possible to significantly refine the algorithms presented here. Neither is it clear that Kraken is necessarily the best option. We chose Kraken as the base of our tool because it is fast and sensitive if a prone to false positives. Other marker based tools may show more or less improvement with the algorithms presented here.

Finally, the algorithms presented here act as a layer on top of a marker based classification tool. This takes advantage of a practical property of Linked-Reads: they can be used as-is with existing tools for short reads. But it potentially leaves room for performance gains. A purpose built Linked-Read taxonomic classifier could search for a parsimonious set of taxonomic assignments for each read cloud.

## Data Access

All raw sequencing reads from this study have been submitted to the NCBI Sequencing Read Archive (SRA; https://www.ncbi.nlm.nih.gov/sra) under accession number PRJNA505182. All code from this study is available in the supplementary materials and at https://github.com/dcdanko/kraken-linked. During review the GitHub link will be private but reviewers are welcome to request access; the exact code used for the paper is available in the supplement. The database used for testing is available for download with the link provided in the supplement.

## Acknowledgement

We sincerely thank 10x Genomics Inc. for providing us valuable real sequencing data sets used in this study for evaluations. We also thank Stephen Williams of 10x Genomics Inc. for coordinating the 10x Metagenomics consortium (a small group of laboratories committed to studying the role of Linked-Reads in metagenomics) in which our team participates. We thank D. Wood, F. Breitwieser and all other authors of Kraken and KrakenUniq for their original work and support.

DCD and DM were supported by the Tri-Institutional Training Program in Computational Biology and Medicine (CBM) funded by the NIH grant 1T32GM083937. This work was also supported by start-up funds (Weill Cornell Medicine) to IH and a National Science Foundation (NSF) grant under award number IIS-1840275 to IH and CM.

